## Supplemental for "FORCE trained spiking networks do not benefit from faster learning while parameter matched rate networks do"

### S3 Fig Methods

Each network used the same randomly initialized decoder, $\boldsymbol{\phi}^{rand}$, sampled from a normal distribution $\mathcal{N}\left(0, N^{-1}\right)$ with zero mean and standard deviation $N^{-1}$. To ensure an exactly zero mean, the sample mean was subtracted from the weights. This was done by first sampling $\tilde{\boldsymbol{\phi}}_{rand} \sim \mathcal{N}\left(0, N^{-1}\right)$ and then defining:

$$\phi_i^{rand} = \tilde{\phi}_i^{rand} - \frac{1}{N}\sum_{j=1}^{N} \tilde{\phi}_j^{rand}. \tag{1}$$

The driving input $y(t)$ was introduced through an input weight vector $\boldsymbol{\psi} \in \mathbb{R}^N$, scaled by the feedback strength parameter $Q$. The input current the $i^{th}$ neuron was thus defined by:

$$I_i^{S/R}(t) = \sum_{j=1}^{N} \omega_{ij}^{S/R} r_j^{S/R}(t) + I_{bias} + Q\psi_i y(t). \tag{2}$$

Each network used the same $\boldsymbol{\psi}$, sampled from a uniform distribution over the interval $[-1, 1]$. The feedback and reservoir weights were generated as described in the Network Models section of the results.

The neural basis correlation was approximated by taking a random sample of 20 neurons, computing the cross-network Pearson correlation, and then taking the average across the 20 samples:

$$\rho\left(\boldsymbol{r}^S, \boldsymbol{r}^R\right) = \frac{1}{20}\sum_{i=1}^{20} \rho(\boldsymbol{r}_i^S, \boldsymbol{r}_i^R), \tag{3}$$

where $\rho$ is the Pearson correlation. The matrices $\boldsymbol{r}^S$ and $\boldsymbol{r}^R$, are the time-sampled neural bases from the LIF and LIF-matched rate networks respectively. The readout correlation was computed by the Pearson correlation between the time-sampled time-series of decoded outputs, $\hat{x}^S(t)$ and $\hat{x}^R(t)$, from the two networks.

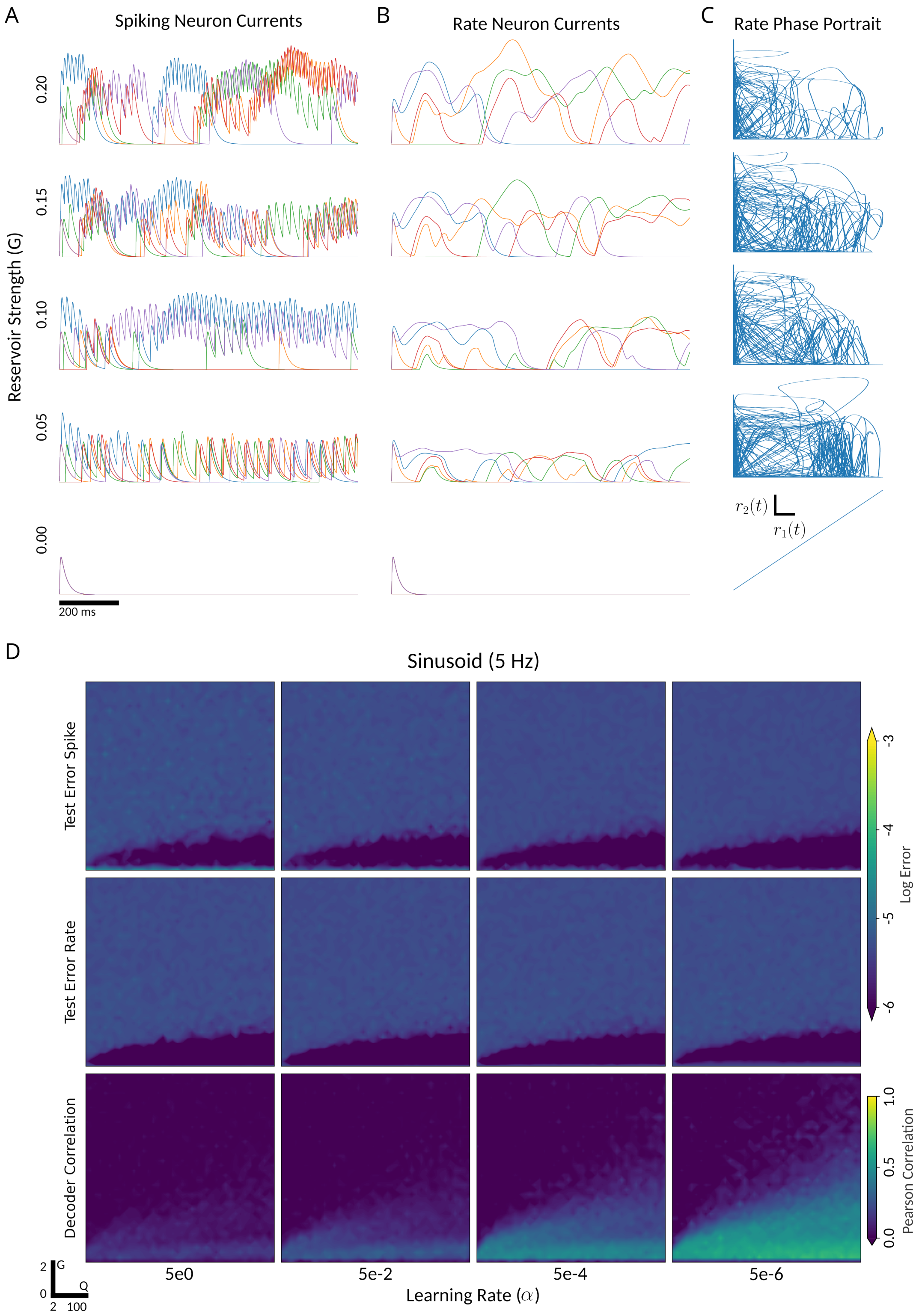


Figure 1: **Chaotic regime for FORCE-trained LIF and LIF-matched rate networks. A-B** Neural currents for networks of 200 LIF neurons and their corresponding LIF-matched rate neurons, demonstrated chaotic behaviour for reservoir strength parameter $G > 0$. **C** Phase portrait of the first two neurons in the rate network simulated for 50 s, displaying chaotic behaviour for $G > 0$. **D** Networks of 2000 LIF and LIF-matched rate neurons were trained over a $40 \times 40 (Q, G)$ parameter grid with $G \in [0, 2]$ and $Q \in [2, 100]$. For sufficiently large $G$, neither network was able to learn.

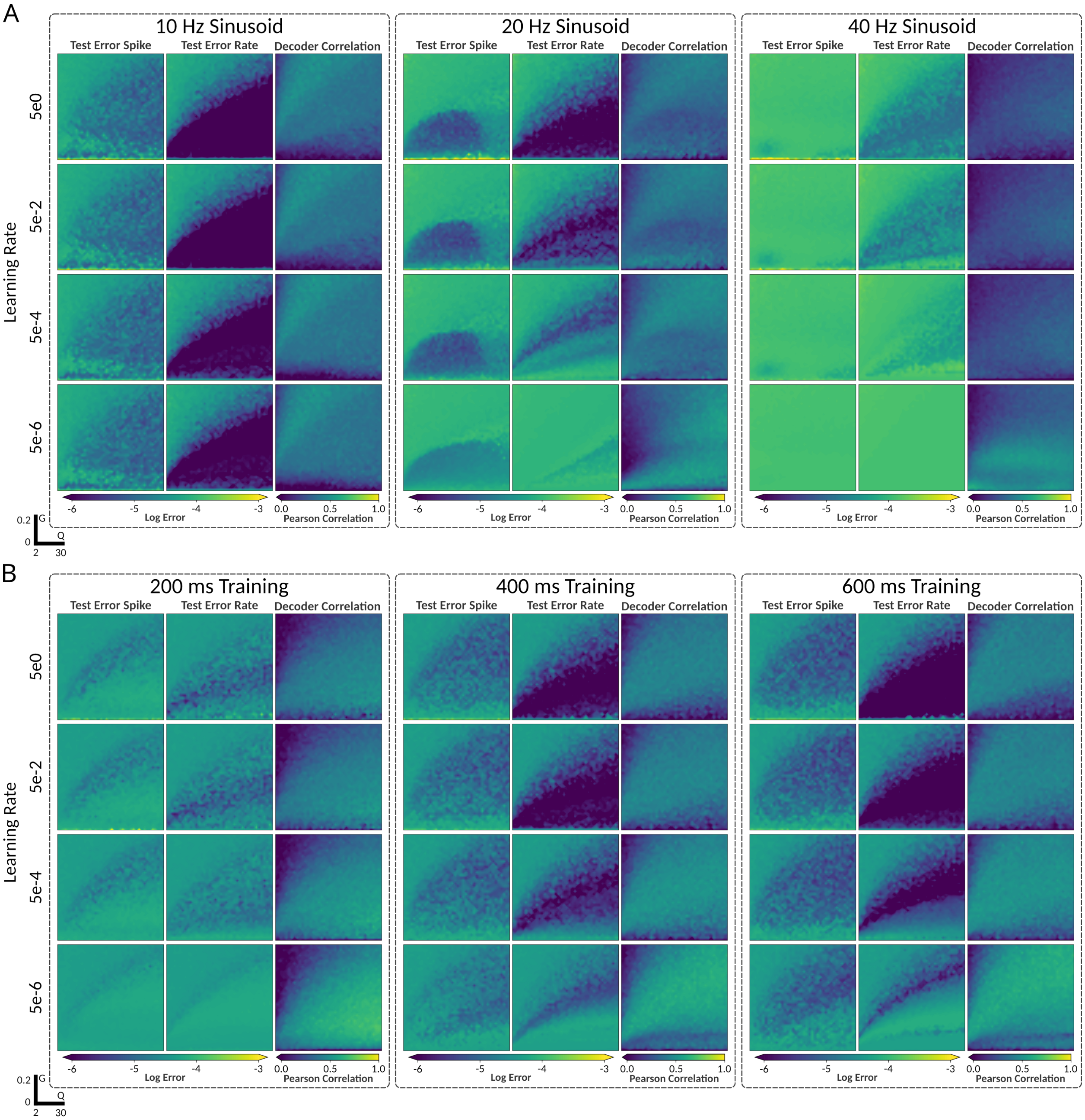


Figure 2: **FORCE-trained networks struggle with fast supervisors and short training durations.** **A** Networks of 2000 LIF and LIF-matched rate neurons were trained to generate sine waves of increasing frequency. As the frequency increased, both networks exhibited reduced ability to learn the supervisor. **B** Networks of 2000 LIF and LIF-matched rate neurons were trained to generate a 5 Hz sine wave with varying training durations. When the training period was very short (containing only a single cycle of the supervisor), neither network was able to learn.

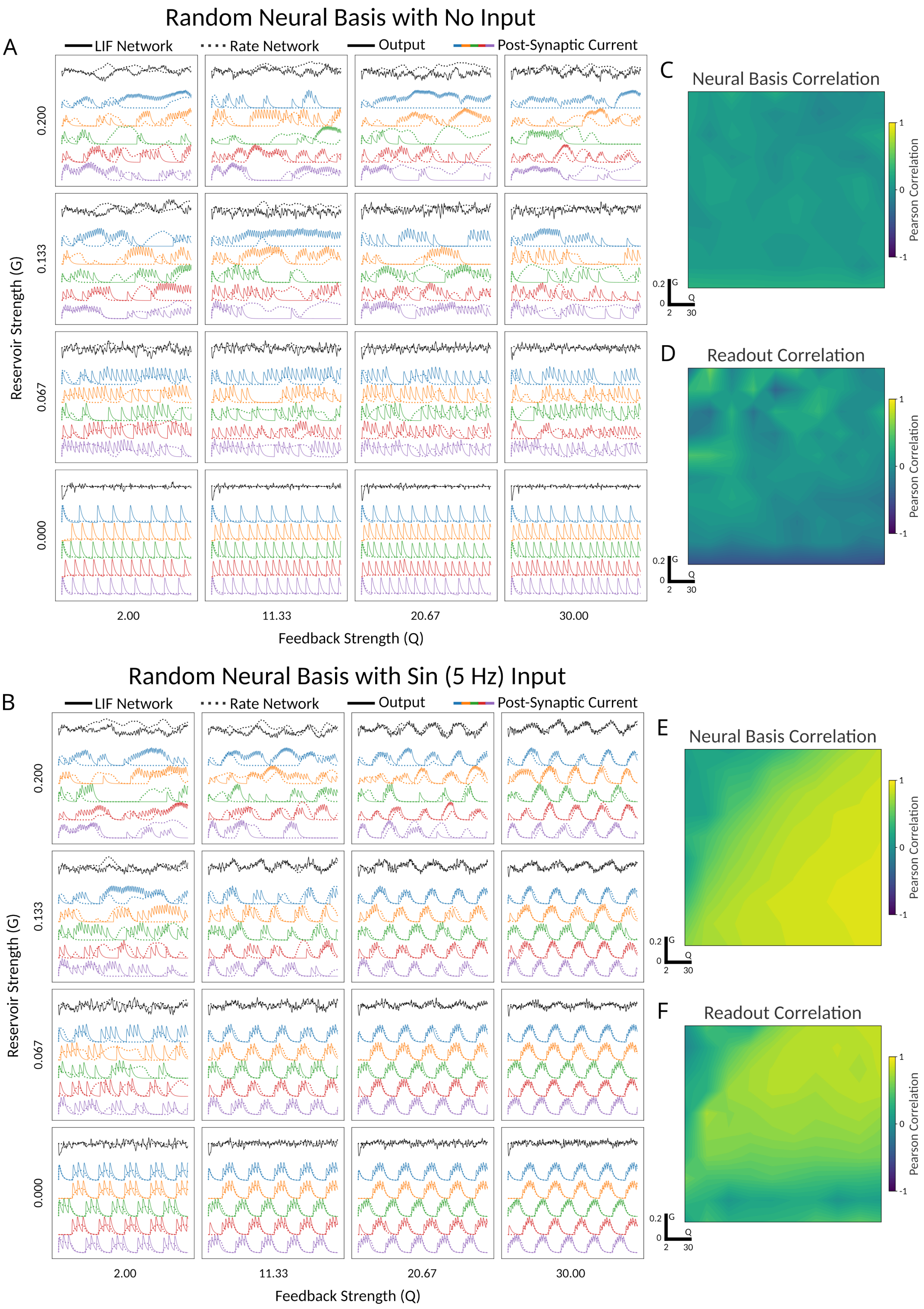


Figure 3: **LIF and LIF-matched rate networks exhibit correlated neural bases when driven but are chaotic without input.** Networks of 2000 LIF and LIF-matched rate neurons were simulated for 1s across different reservoir strengths ($G$) and feedback strengths ($Q$), both with and without driving input. In the absence of input, both networks exhibited chaotic dynamics and low cross-network correlations. When driven, the cross-network readout and neural bases became correlated. **A-B** Sample readouts and neural basis elements from both networks. **C-F** Cross-network correlations of sampled neural bases and readouts.

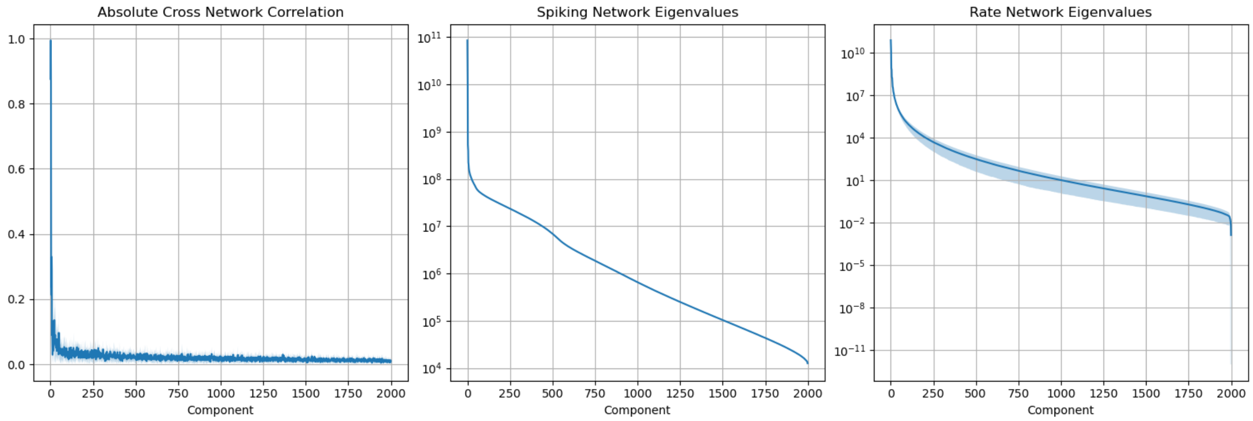


Figure 4: **LIF and LIF-matched rate networks have highly correlated low order principle components.** Networks of 2000 LIF and LIF-matched rate networks were trained with FORCE on the 5 Hz sinusoidal supervisor for 4s with learning rate $\alpha$ =5e-6, reservoir strengths $G = 0.1$, and feedback strengths $Q = 15$, for 10 different seed values. We then plot the averaged: absolute correlation in the orthogonal basis elements, LIF eigenvalue, and Rate eigenvalue. The shaded regions represent the standard deviation. The lower order orthogonal basis elements have much higher associated eigenvalues and so explain much more of the variability in the original basis. The early basis elements are also highly correlated across networks types.
